# The Deubiquitinase USP36 Regulates Growth and Spermatogenesis Through Catalytic-Dependent and -Independent Mechanisms in *Drosophila*

**DOI:** 10.1101/2025.03.25.644999

**Authors:** Carmen Coirry, Julie Manessier, Charlène Clot, Magda Mortier, Marie-Odile Fauvarque, Emmanuel Taillebourg

## Abstract

Deubiquitinases (DUBs) form a specific class of proteases removing ubiquitin from target proteins. They are involved in the regulation of many cellular processes including cell growth and proliferation. Among them, USP36 is a key regulator of the oncogenic transcription factor c-Myc, preventing its degradation by the proteasome. These two proteins form an evolutionary conserved complex providing the opportunity to investigate USP36 mechanisms of action *in vivo* in a genetically tractable model such as *Drosophila melanogaster*. Null mutants of *dUsp36* die early during larval development and exhibit severe growth defects. Strikingly, we report here that flies carrying a CRISPR/Cas9-induced catalytic mutation of *dUsp36* survive to adulthood with only minor growth defects, yet males are infertile. This finding indicates that dUSP36 deubiquitinating activity is dispensable for cell growth but essential for spermatogenesis. Our results thus reveal that dUSP36 functions through both catalytic-dependent and catalytic-independent mechanisms, highlighting a dual mode of action with implications for the understanding of DUBs mechanism of action.

## INTRODUCTION

Deubiquitinases (DUBs) are proteases that specifically remove ubiquitin moieties from ubiquitinated proteins. They regulate numerous cellular processes (Komander et al. 2009; Clague et al. 2013; Leznicki and Kulathu 2017) and their dysregulation has been linked to numerous human cancers (Sacco et al. 2010; Fraile et al. 2018; Dewson et al. 2023). In particular, USP36, a DUB primarily localized in the nucleolus, supports ribosomal biogenesis and maintains nucleolar integrity by deubiquitinating and stabilizing multiple nucleolar proteins, such as c-MYC, DHX33, SNAIL1, and RNA Pol I (Endo, Kitamura, et al. 2009; Endo, Matsumoto, et al. 2009; Sun, He, et al. 2015; Fraile et al. 2018; van den Heuvel et al. 2021; Qin et al. 2023; Yang et al. 2024). The role of USP36 in c-MYC regulation has been extensively studied (Sun, He, et al. 2015; Sun, Sears, et al. 2015; Liu et al. 2019; Thevenon et al. 2020; Hu and Luo 2024), as precise control of c-MYC levels is critical for normal cell growth and proliferation, while its dysregulation contributes to tumorigenesis (van Riggelen et al. 2010; Miller et al. 2012; Sabo and Amati 2014; Grifoni and Bellosta 2015).

We previously showed that in *Drosophila melanogaster*, dUSP36 is essential for cell growth, and stabilizes dMYC (Thevenon et al. 2020). To further investigate the *in vivo* mechanisms of action of dUSP36 and assess the specific role of its DUB activity, we generated a catalytically inactive version of the endogenous dUSP36 protein using CRISPR/Cas9 mutagenesis. Here, we demonstrate that unlike *dUsp36* null mutants, which completely lack dUSP36 protein and exhibit severe growth defects leading to larval lethality, *dUsp36* catalytic mutants display only minor growth defects and develop to adulthood. Interestingly, *dUsp36* catalytic mutant males exhibit severely reduced fertility.

Altogether, these findings reveal that while the presence of the dUSP36 protein is essential for cellular growth, its catalytic activity is dispensable for this process but critically required for spermatogenesis. These results demonstrate that, *in vivo*, dUSP36 functions through both catalytic-dependent and catalytic-independent mechanisms, revealing a dual mode of action.

## RESULTS

### Generation of *dUsp36* catalytic mutants by CRISPR/Cas9 mutagenesis

To generate a catalytically inactive version of the dUSP36 protein, we mutated the cysteine residue (C181) within a conserved domain previously shown to be essential for DUB catalytic activity (Hu et al. 2005; Komander et al. 2009; Faesen et al. 2011; Ye et al. 2011) to a serine residue (Fig. S1A; see Materials and Methods for details). This C181S mutation has been previously demonstrated to completely abolish dUSP36 catalytic activity (Buszczak et al. 2009). The CRISPR/Cas9 mutagenesis yielded two independent *dUsp36*^*C181S*^ mutant strains, as well as a control strain, *dUsp36*^*CTL*^, which contains the same silent mutations introduced for screening purposes as the mutant alleles but lacks the G-to-C transversion encoding the C181S mutation (Fig. S1C).

Western blot analysis of dUSP36 expression revealed that the dUSP36^C181S^ protein is detected at lower levels than the wild-type dUSP36 protein, whereas the *dUsp36*^*CTL*^ allele produces normal dUSP36 protein levels (Fig. 1A, B). This observation aligns with previous findings indicating that dUSP36 undergoes auto-deubiquitination to regulate its stability (Thevenon et al. 2020). To ensure comparable dUSP36 protein levels across genotypes, we compared *dUsp36*^C181S^ mutants with *dUsp36*^*CTL*^/*dUsp36*^*Δ43*^ heterozygous controls (Fig. 1B). Importantly, the subcellular localization of the dUSP36^C181S^ protein remains indistinguishable from that of the wild-type dUSP36 protein (Fig. 1C). These data confirm the successful generation of the catalytically inactive dUsp36^C181S^ mutant allele, providing a valuable tool for investigating the specific role of dUSP36 catalytic activity *in vivo*.

**Figure 1:**
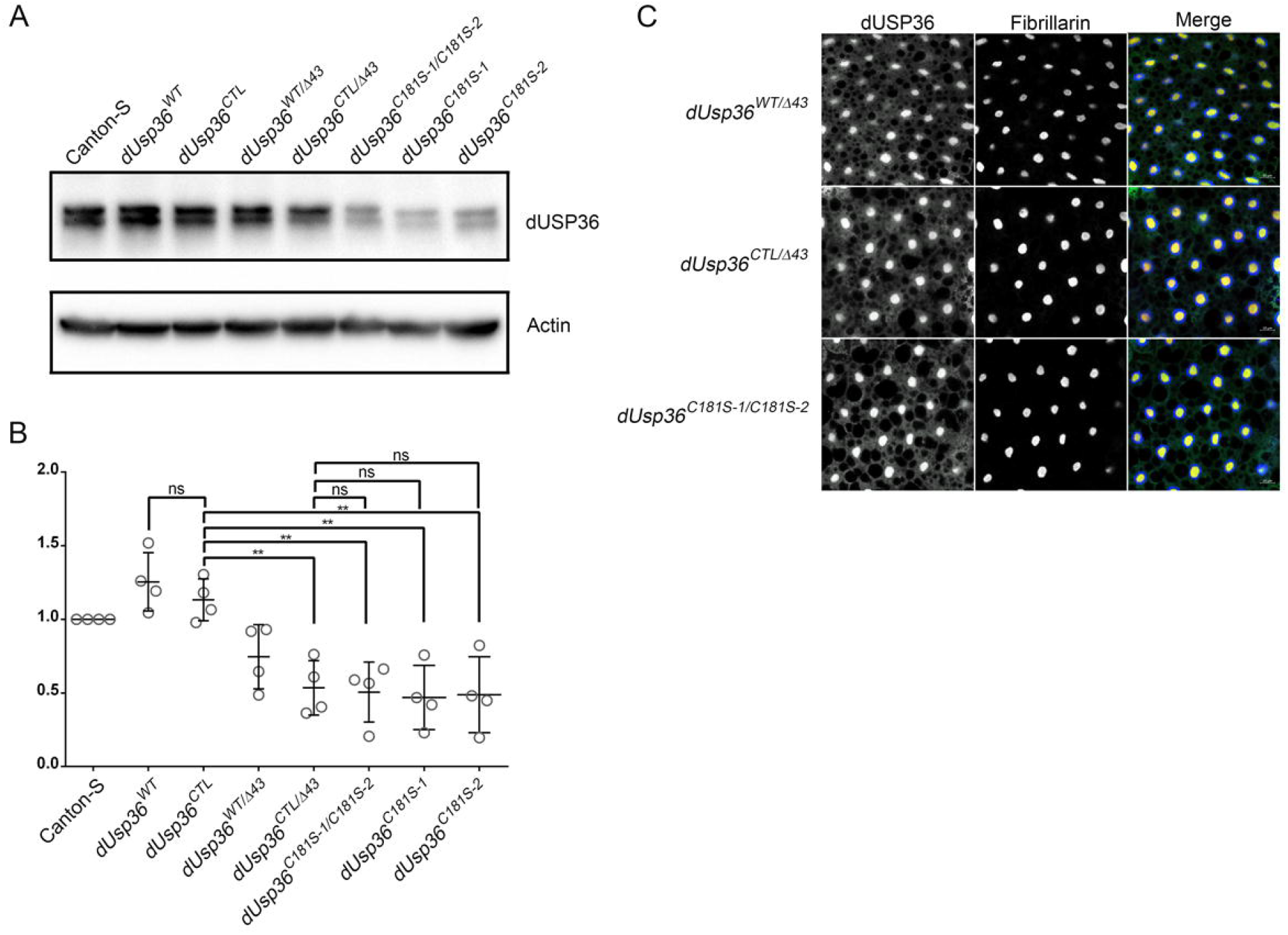
Expression of the dUSP36^C181S^ protein. (A) Total protein extracts from *Drosophila* males of the indicated genotypes were analysed by Western blot using a specific anti-dUSP36 antibody. (B) Quantification of dUSP36 relative levels, expressed as the ratio of dUSP36 signal intensity to Actin signal intensity, normalized to the wild-type Canton-S strain. N = 4. ** indicates a statistically significant difference (P < 0.01) as determined by one-way ANOVA. (C) Fat bodies of third-instar larvae stained with anti-dUSP36 and anti-Fibrillarin antibodies. Images are shown as separate grayscale channels and as merged images. Blue: DAPI. Green: anti-dUSP36. Red: anti-Fibrillarin.

### *dUsp36* catalytic mutants are viable and exhibit a subtle growth phenotype

We previously reported that *dUsp36*^*Δ*43^ null mutants, which completely lack dUSP36 protein, exhibit severe cell growth defects and early larval lethality (Taillebourg et al. 2012) indicating that the presence of dUSP36 protein is required for cellular and organismal growth. In sharp contrast with these previous findings, we found that *dUsp36*^*C181S*^ mutants develop to adulthood without lethality (Fig 2A) and exhibit only a minor reduction in body weight (Fig. 2B), along with a slight decrease in wing size (Fig. 2C, D).

**Figure 2:**
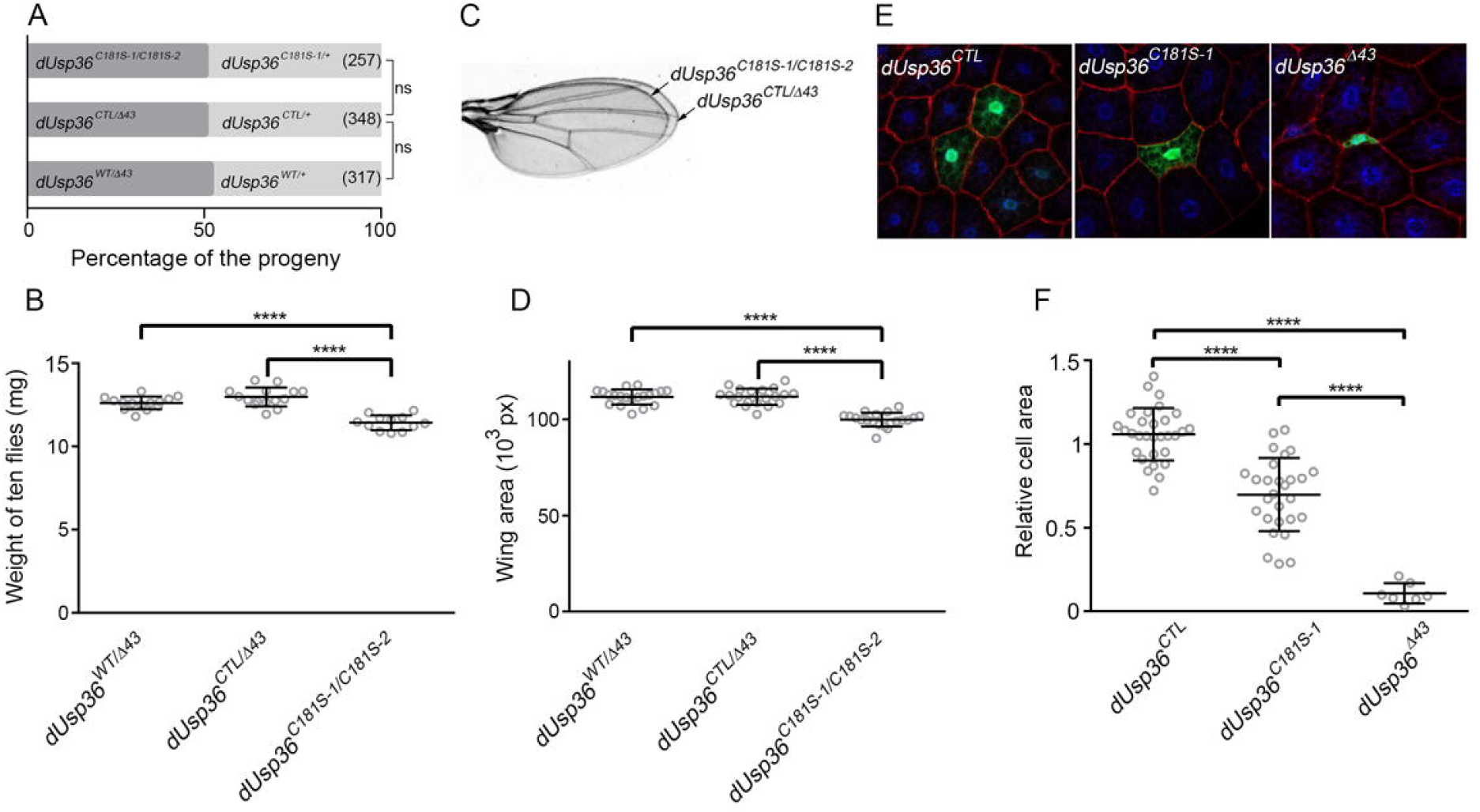
Phenotype of *dUsp36*^*C181S*^ mutants. (A) Histograms showing the observed frequency of the indicated genotypes in the progeny resulting from the following crosses: *dUsp36* ^*WT*^ x *dUsp36Δ*^43^/*TM6b, dUsp36*^*CTL*^ x *dUsp36Δ*^43^/*TM6b*, and *dUsp36*^C181S-1^ x *dUsp36*^C181S^-2/*TM6b* respectively. (B) Weight of groups of ten *Drosophila* females of the indicated genotype. (C) Dissected wings and (D) wing area quantifications. (E) GFP-positive clones of larval fat body cells homozygous for the indicated *dUsp36* allele. (F) Quantifications of the relative cell size, expressed as the ratio between the area of a GFP+ cell and the mean area of neighbouring GFP-cells (F). Genotype: *hsFLP*/+; *CgGal4, UAS-GFP*/+; *FRT2A dUsp36* ^x^/*FRT2A TubGal80*. **** and *** indicate statistical significance with P < 0.001 and P < 0.01, respectively, as determined by chi-square test (A) and one-way ANOVA (B, D, F).

The enzymatic activity of USPs relies on a catalytic triad—a conserved set of three amino acids consisting of a cysteine, a histidine, and an aspartate or asparagine. Histidine-to-aspartate substitutions have also been reported to render the enzyme catalytically inactive (Kovalenko et al. 2003; Hu et al. 2005; Komander et al. 2009). We mutated both the cysteine and histidine residues and generated transgenic flies expressing the dUSP36^C181S, H439D^ protein. When ubiquitously expressed in a *dUsp36* null mutant background, this double mutant protein rescues the larval lethality of *dUsp36* null mutants and allows viable adults to emerge (Fig. S2). These results confirm that the catalytic activity of dUSP36 is not essential for cellular growth and organismal viability.

To determine whether the observed phenotypes result from cell-autonomous factors, we generated clones of cells homozygous for either the control allele *dUsp36* ^*CTL*^, the catalytically inactive allele *dUsp36*^C181S^, or the null allele *dUsp36*^*Δ43*^ (Fig. 2E). This analysis revealed that *dUsp36*^*C181S*^ mutant cells are significantly smaller than neighboring wild-type cells, although their growth is much less affected than that of null *dUsp36* mutant cells (Fig. 2F). This observation indicates that the subtle growth defect of *dUsp36* catalytic mutants is cell-autonomous.

Altogether, these results demonstrate that while the presence of dUSP36 protein is essential for cellular and organismal growth, its catalytic activity plays only a minor, cell-autonomous role in this process.

### Altering dMYC protein levels in *dUsp36* catalytic mutants

Experimental data obtained in human cellular models (Kovalenko et al. 2003; Hu et al. 2005; Komander et al. 2009) provide compelling evidence that USP36 depletion affects the stability of nucleolar c-MYC and inhibits cell proliferation. To test whether the minor growth defects of *dUsp36* catalytic mutants depend on dMYC function, we evaluated the phenotypic consequences of altered dMYC protein levels in *dUsp36* wild-type or mutant contexts. To increase dMYC levels, we introduced a transgene expressing a GFP-tagged version of dMYC within its genomic environment, providing additional *dMyc* copies alongside the wild-type *dMyc* locus (Greer et al. 2013). This transgene fully rescues the lethality of *dMyc*^*4*^ null mutants, demonstrating that it expresses physiological levels of functional dMYC protein (data not shown). Remarkably, this transgene significantly enhances the body weight of control flies but not that of *dUsp36* catalytic mutants (Fig. 3A). These results indicate that while control flies respond to dMYC overexpression, *dUsp36* catalytic mutants do not. To further explore the genetic interaction between *dMyc* and *dUsp36*, we used the *dMyc*^*P0*^ hypomorphic mutation that decreases dMYC protein levels and leads to reduced body weight (Johnston et al. 1999). *dUsp36* catalytic mutants and *dMyc*^*P0*^ mutants develop to adulthood without lethality (Fig. 3B). In contrast, in a *dMyc*^*P0*^ genetic background, the frequency of *dUsp36* mutants is significantly reduced (Fig. 3B), whereas no such reduction of viability is observed in *dUsp36*^*C181S*^ control flies. This implies that while dUSP36 catalytic activity is dispensable under normal conditions, it becomes critical when *dMyc* function is compromised.

**Figure 3:**
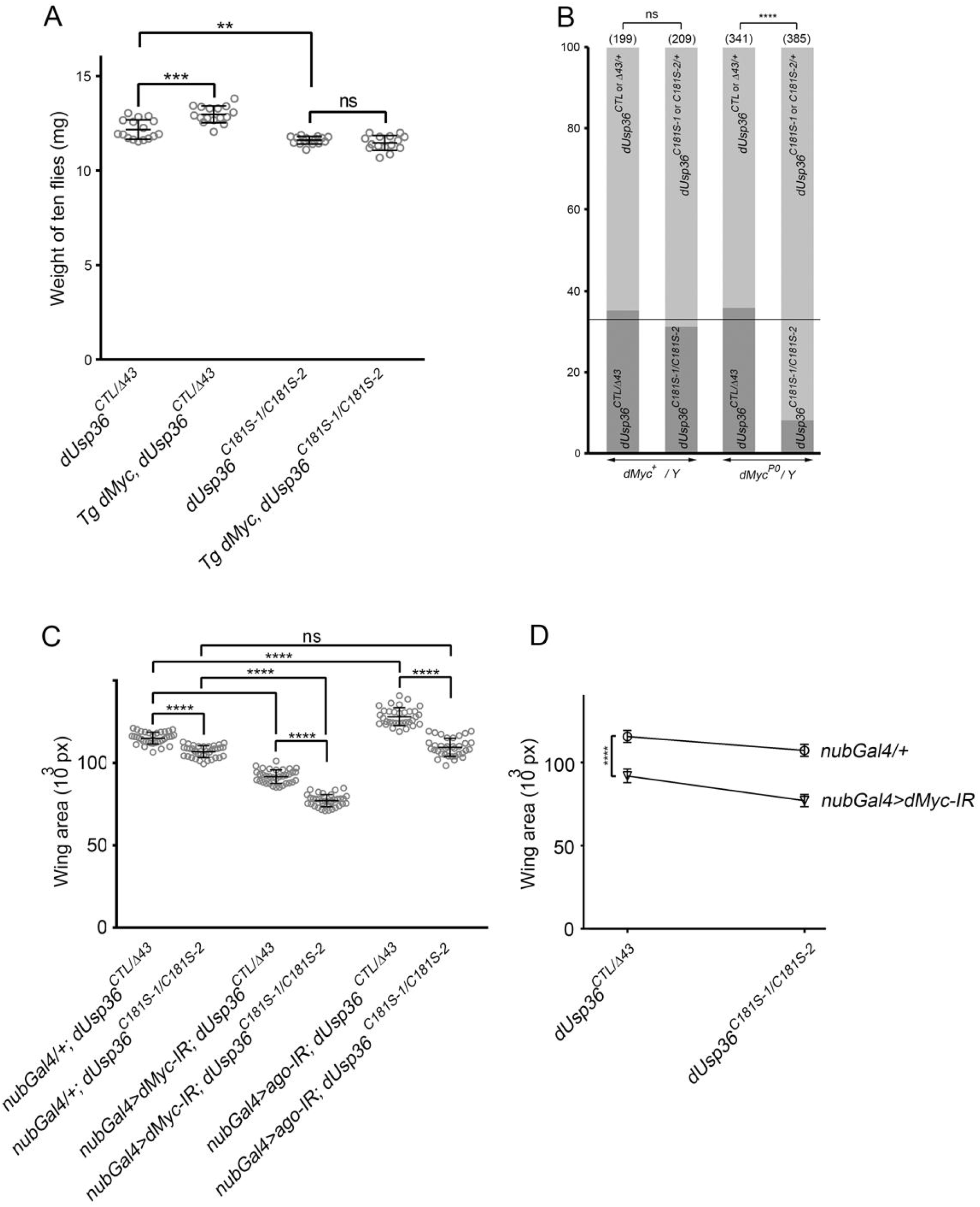
Phenotypic effects of dMYC levels increase and decrease in *dUsp36* catalytic mutants. (A) Quantifications and statistical analyses of female fly weight for the indicated genotypes. “*Tg dMyc*” indicates the presence at the homozygous state of a transgene expressing a GFP-tagged version of dMYC within its genomic environment. (B) Histograms showing the observed frequency of males of the indicated genotypes in the progeny resulting from the following crosses: *dUsp36*^*CTL*^/*TM6b* x *dUsp36Δ*^43^/*TM6b, dUsp36*^C181S-1^/*TM6b* x *dUsp36*^C181S^-^2^/*TM6b, dMyc*^P0^/*FM7c*; *dUsp36*^*CTL*^/*TM6b* x *dUsp36*^*Δ43*^/*TM6b*, and *dMyc*^P0^/*FM7c*; *dUsp36*^C181S-1^/*TM6b* x *dUsp36*^C181S^-2/*TM6b*, respectively. (C, D) Quantifications and statistical analyses of wing areas of control and catalytic mutant flies expressing the wing-specific Gal4 driver *nubGal4*, either alone or in combination with RNAi transgenes targeting *dMyc* or *ago*. **** and *** indicate statistical significance with P < 0.001 and P < 0.01, respectively, as determined by one-way ANOVA (A, C), chi-square test (B), and two-way ANOVA (D).

To determine whether these findings hold true at the organ level, we compared the effects of altered dMYC protein levels in the developing wing between control flies and *dUsp36* catalytic mutants (Fig. 3C). In this setup, wing-specific increases of dMYC protein levels were achieved by reducing AGO protein levels using a specific RNAi transgene targeting *Ago*. Indeed, AGO is the F-box component of the SCF E3 ligase complex, which ubiquitinates dMYC and negatively regulates its protein levels *in vivo*. Knocking down *Ago* leads to elevated dMYC protein levels and increased tissue growth (Moberg et al. 2004). As expected, in control flies, reducing AGO protein levels results in increased wing area (Fig. 3C). In contrast, in the developing wing of *dUsp36* catalytic mutants, it does not significantly increase wing size (Fig. 3C). These results indicate that increased dMYC levels promote wing growth in control flies but not in *dUsp36* catalytic mutant flies.

Conversely, dMYC protein levels were specifically reduced by expressing a *dMyc* targeting RNAi transgene in the developing wing. In that context, both control flies and *dUsp36* catalytic mutants displayed reduced wing size (Fig. 3C). A two-way ANOVA analysis reveals that *dUsp36* catalytic mutants are more affected by dMYC depletion than control flies (Fig. 3D).

Altogether these results show that increased dMYC levels fail to increase body weight or wing area in *dUsp36* catalytic mutants, whereas they do in control flies. This suggests that reduced dMYC protein levels are not the primary cause of the growth defect of *dUsp36* catalytic mutants. In contrast, *dUsp36* catalytic mutants exhibit a strong sensitivity to dMYC depletion, suggesting that, *in vivo*, dUSP36 catalytic activity acts in the same pathway as dMYC to promote growth.

### *dUsp36* catalytic mutation affects male fertility

Specific combinations of viable *dUsp36* loss-of-function alleles have been shown to affect male and female fertility (Buszczak et al. 2009). Here, we observed here that *dUsp36*^*C181S*^ females are slightly, but significantly, less fertile than control females (Fig. 4A). It remains unclear however whether this decreased fertility is due to a general effect related to their reduced body weight or a specific role of dUSP36 catalytic activity in female gametogenesis. Strikingly, *dUsp36* catalytic mutant males produce significantly fewer offspring than control males, with most being completely sterile (Fig. 4B). Consistent with this observation, *dUsp36*^*C181S*^ males exhibit reduced seminal vesicle size, indicating lower sperm production compared to controls. Moreover, we show here that dUSP36 protein is widely expressed during spermatogenesis (Fig. 4C, D). These findings reveal that dUSP36 catalytic activity plays a critical role in spermatogenesis.

**Figure 4:**
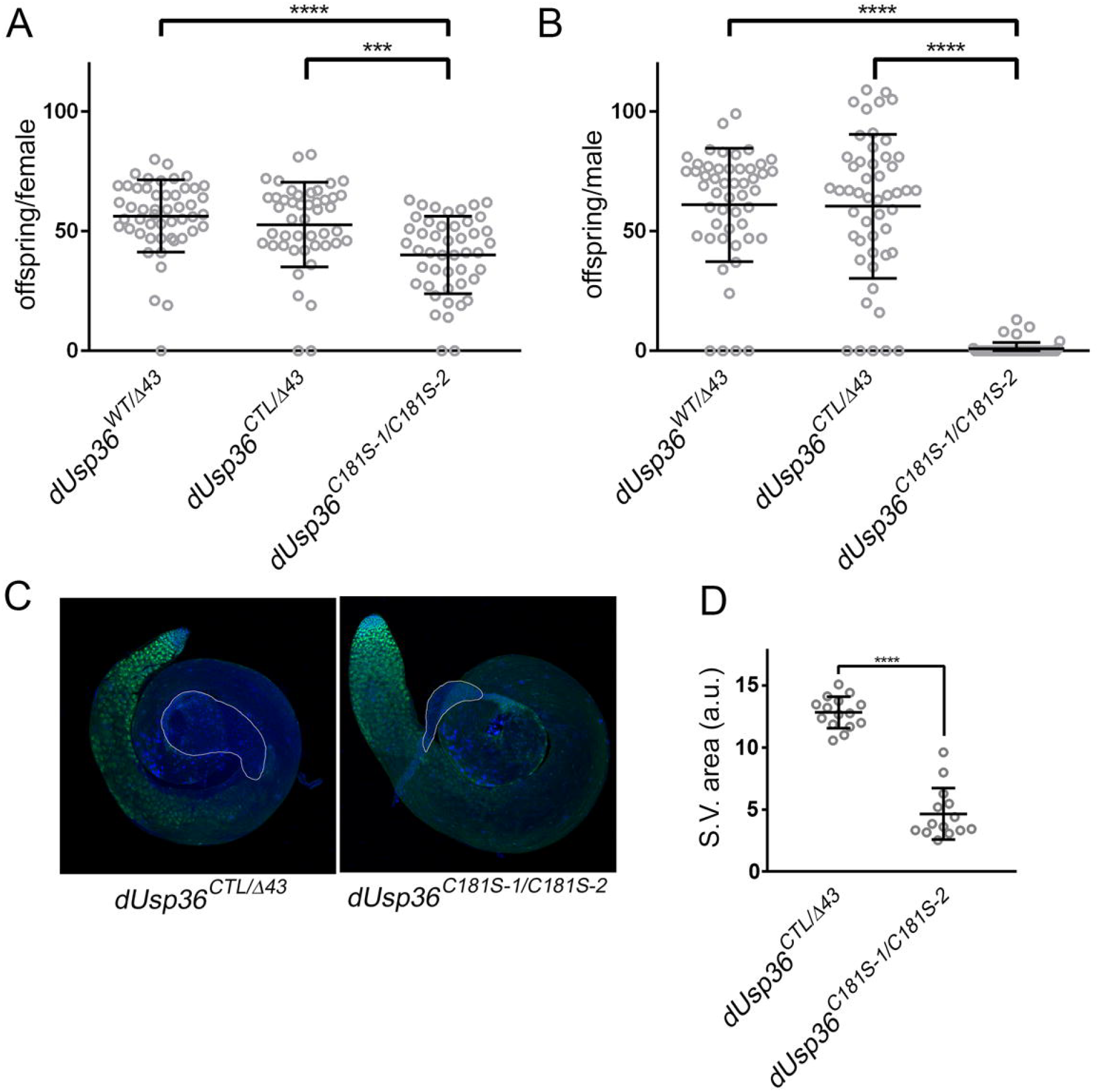
Fertility defects in *dUsp36* catalytic mutants. Quantifications and statistical analyses of the number of offspring produced by individual females (A) and males (B) of the indicated genotypes. (C) Testis stained with anti-dUSP36. The seminal vesicle is outlined. Blue: DAPI. Green: anti-dUSP36 antibody. (D) Quantifications and statistical analyses of the area of the seminal vesicle (S.V.). **** and *** indicate statistical significance with P < 0.001 and P < 0.01, respectively, as determined by one-way ANOVA.”**

## DISCUSSION

The data presented here demonstrate that inactivating the catalytic activity of the DUB USP36 in *Drosophila* is not equivalent to its complete or partial depletion. In particular, the catalytic mutant does not display the cellular and organismal growth defects observed in null *dUsp36* mutant. This indicates that dUSP36 functions primarily through non-catalytic mechanisms in these processes. In contrast we report that USP36 DUB activity is essential for male fertility, making it the only process identified so far that strictly depends on USP36 enzymatic function.

It is unlikely that the *dUsp36*^*C181S*^ mutation fails to fully inhibit the DUB activity *in vivo*. Indeed, the lower protein levels of dUSP36^C181S^ compared to wild-type dUSP36 (Fig. 1A, B) indicate that it fails to stabilize itself, unlike the wild-type form which undergoes auto-deubiquitination, strongly suggesting that this protein is indeed catalytically inactive (Thevenon et al. 2020). Furthermore, a double-mutant dUSP36 protein, in which both the catalytic cysteine and the histidine residue of the conserved catalytic triad are mutated, is still able to sustain cell and organismal growth in a null mutant background *in vivo*. These results strongly argue against the hypothesis that the minor growth defect observed in dUsp36^C181S^ mutants is due to residual DUB activity.

We further demonstrate that *dUsp36* catalytic mutants do not respond to increased dMYC levels. This suggests that reduced dMYC levels do not contribute to the minor growth defects observed in *dUsp36*^*C181S*^ mutants. However, these mutants are highly sensitive to dMYC depletion, indicating that dUSP36 catalytic activity likely plays a role in the same pathway as dMYC. Taken together, our results indicate that dUSP36 primarily functions independently of its enzymatic activity in promoting cell growth and that – in contrast with previous reports - its catalytic activity is not required for dMYC *stabilization in vivo*.

These results should be considered in the context of existing literature, including our own work, showing that in *Drosophila* and human cultured cells, USP36 deubiquitinates and stabilizes MYC (Sun, Sears, et al. 2015; Sun, He, et al. 2015; Thevenon et al. 2020). Several hypotheses could explain this discrepancy: (1) The deubiquitination and stabilization of MYC by USP36 observed in cultured cells may also occur *in vivo* but could be compensated for by redundant regulatory mechanisms. If this is the case, USP36 would be expected to regulate another dMYC-independent growth process through a non-catalytic mechanism. Along these lines, recent studies in human cells indicate that USP36 functions as a SUMO ligase to promote the SUMOylation of specific nucleolar proteins involved in ribosome biogenesis, a process that is crucial for cell and organism growth (Ryu et al. 2021; Chen et al. 2023; Li et al. 2023; Li et al. 2024; Yang et al. 2024). In human cells, the catalytic cysteine of USP36 is essential for both its DUB and SUMO ligase activities (Ryu et al. 2021). If *Drosophila* dUSP36 functions similarly as its human orthologue, *dUsp36*^*C181S*^ mutants should also lack SUMO ligase activity, which would impair ribosome biogenesis and induce severe growth defects similar to those observed in *Minute* mutants (Marygold et al. 2007). However, the mild growth phenotype of *dUsp36* ^*C181S*^ mutants does not support this hypothesis. Additionally, we failed to detect SUMO ligase activity for dUSP36 in *Drosophila* cells (data not shown). (2) Contrary to previous conclusions based on knock-down experiments, USP36 may control MYC stability independently of its catalytic activity: it may not directly deubiquitinate MYC but instead regulate its ubiquitination level indirectly. This could occur through inhibition of an E3 ligase targeting MYC or recruitment of another DUB within the regulatory complex. In this model, USP36 knock-down via RNAi or loss-of-function mutations would impair these functions, leading to MYC destabilization. Moreover, overexpressing wild-type USP36 may result in non-specific deubiquitination of MYC, whereas a catalytically inactive USP36 mutant would not act ectopically, falsely leading to the conclusion that USP36 directly deubiquitinates MYC (Sun, He, et al. 2015; Thevenon et al. 2020). Additional experiments will be needed to clarify the exact mechanisms by which USP36 regulates MYC and contributes to cell growth.

While dUSP36 catalytic activity plays a minor role in cell and organismal growth, our results clearly demonstrate that it is essential for spermatogenesis. This is consistent with previous findings showing that dUSP36 is required for germline stem cell maintenance by deubiquitinating histone H2B and repressing premature expression of differentiation genes (Buszczak et al. 2009).

In conclusion, our findings indicate that, in *Drosophila*, dUSP36 functions through a dual mechanism: in cell growth, dUSP36 acts independently of its catalytic activity, similar to other DUBs or pseudo-DUBs (Campos Alonso and Knobeloch 2024) whereas in spermatogenesis, its DUB activity is crucial, highlighting a specialized catalytic function in germline development. Our results therefore reshape our understanding of how DUBs functions during cellular responses and development, extending beyond their enzymatic activity. Determining whether this dual mechanism is unique to dUSP36 or extends to its human counterpart and other DUBs will be of great interest.

## MATERIALS AND METHODS

### Drosophila strains

The Bloomington Drosophila Stock Center provided the following Drosophila strains: *Tg dMyc* ^*P0*^ (#81274), *dMyc* (#11298), *UAS-dMyc-IR* (# 25783), *UAS-Ago-IR* (#34802), *nubGal4* (#25754), *hsFLP* (#8862), *FRT2A Gal80* (#5190), *FRT2A* (#1997) and *CgGal4* (#7011).

### CRISPR/Cas9 mutagenesis

The gRNA sequence was selected using the E-CRISP website and cloned into the pCFD3 plasmid (Port et al. 2014). A 169-nucleotide long single stranded oligonucleotide donor (ssODN) repair template was synthesised (https://eu.idtdna.com/page). It contains the mutation of interest, a silent mutation creatin g a Pst1 restriction site as well as silent mutations in the PAM and the gRNA to avoid re-cleaving of an edited chromosome (Sup. Fig. 1A). The ssODN was injected along with the sgRNA expression vector (ssODN: 100 ng/uL and sgRNA: 250 ng/uL) into the *nos-Cas9* (Port et al. 2014) recipient strain (BestGene Inc.). Sixty-six F0 founders were crossed individually and up to 10 F1 males were crossed individually for each F0 founder; each F1 male was then retrieved for single fly PCR. After amplification, PCR products were digested by Pst1, leading to the identification of six independent Pst1 positive strains (Sup. Fig. 1B). Sequencing revealed that three of them contained the desired C181S mutation (two, *dUsp36*^*C181S-1*^ and *dUsp36*^*C181S-2*^, were kept for further analysis) whereas three others did not include the C181S mutation but contains all the silent mutations (Sup. Fig. 1C). One of them, *dUsp36* ^*CTL*^, was used as control.

To ensure that the mutagenesis process did not add any off-target mutations, the genomic region was sequenced and these alleles were outcrossed for six generations on a wild-type chromosome. Moreover, all the phenotypic analyses were performed on heterozygotes (*dUsp36*^*CTL*^/*dUsp36*^*Δ43*^ and *dUsp36*^*C181S-1*^/*dUsp36*^*C181S-2*^).

### Western blots

Protein lysates were separated on SDS-PAGE gels (TGX Stain Free, BioRad). Proteins were transferred to PVDF membranes (BioRad) which were treated as already described (Thevenon et al. 2020). Image acquisitions were performed with the Chemidoc imaging system (BioRad) and quantifications using the Image Lab software (BioRad). The anti-dUSP36 antibody, a kind gift from Dr. M. Buszczak, was used at a 1/2500 dilution

### Immunohistochemistry

Lateral lobes of third instar larval fat bodies or adult testes were dissected in PBS, fixed for 30 min in 4% paraformaldehyde and rinsed twice in PBS. The samples were then blocked for 1 h in PBS, 0.1% Triton X-100, 5% normal goat serum and incubated overnight at 4°C with the anti-dUSP36 (1/200) and anti-Fibrillarin 38F3 (1/200, abcam) antibodies. Secondary antibodies were coupled to Alexa594 or Alexa488 (1/500, Invitrogen). After mounting in DAPI containing Vectashield (Vector Laboratories, H-1200), the samples were imaged using a Zeiss LSM 880 confocal laser scanning microscope.

### Phenotypic analysis

For weight measurements, the larval density was carefully controlled. Three-day-old females were weighted by groups of ten on a Mettler Toledo Classic precision balance. Adults and dissected wings were imaged using a Keyence VHX-5000 numerical microscope. Wing areas were blindly measured manually using the Fiji/ImageJ software (National Institute of Health). For clonal analysis of cell size in the fat body, 0–6 h embryos were subjected to heat-shock for 1 h at 37°C and resulting 4.5 days old L3 larvae were dissected. Fat body’s lateral lobes were fixed for 30 min in 4% paraformaldehyde, rinsed twice in PBS, and incubated overnight at 4°C in 2 U/ml Alexa Fluor 546 Phalloidin (Invitrogen, A22283). After mounting in DAPI containing Vectashield (Vector Laboratories, H-1200), the samples were imaged using a Zeiss LSM 880 confocal laser scanning microscope. Cell size was quantified using the Fiji/ImageJ software (National Institute of Health) as the ratio between the area of a GFP+ cell and the mean area of neighbouring GFP-cells.

For fertility tests, female and male flies of the indicated genotype were collected, aged for 3⍰days, and individually mated to wild-type flies. The vials were emptied after a three-day period and the progeny was counted.

Statistical analyses were performed using the GraphPad Prism6 software.

## AKNOWLEDGEMENT

We thank Dr. Buszczak for the anti-dUSP36 antibody, the Bloomington *Drosophila* Stock Center for providing *Drosophila* strains, and the IRIG µLife facility for confocal microscopy. This work was supported by the French National Research Agency in the framework of the “Investissements d’avenir” program (ANR-17-EURE-0003) through a funding from GRAL. Thesis operating costs are supported by IDEX Université Grenoble Alpes and GRAL PhD Operating Costs (CC).

## FIGURE LEGENDS

**Figure S1:**
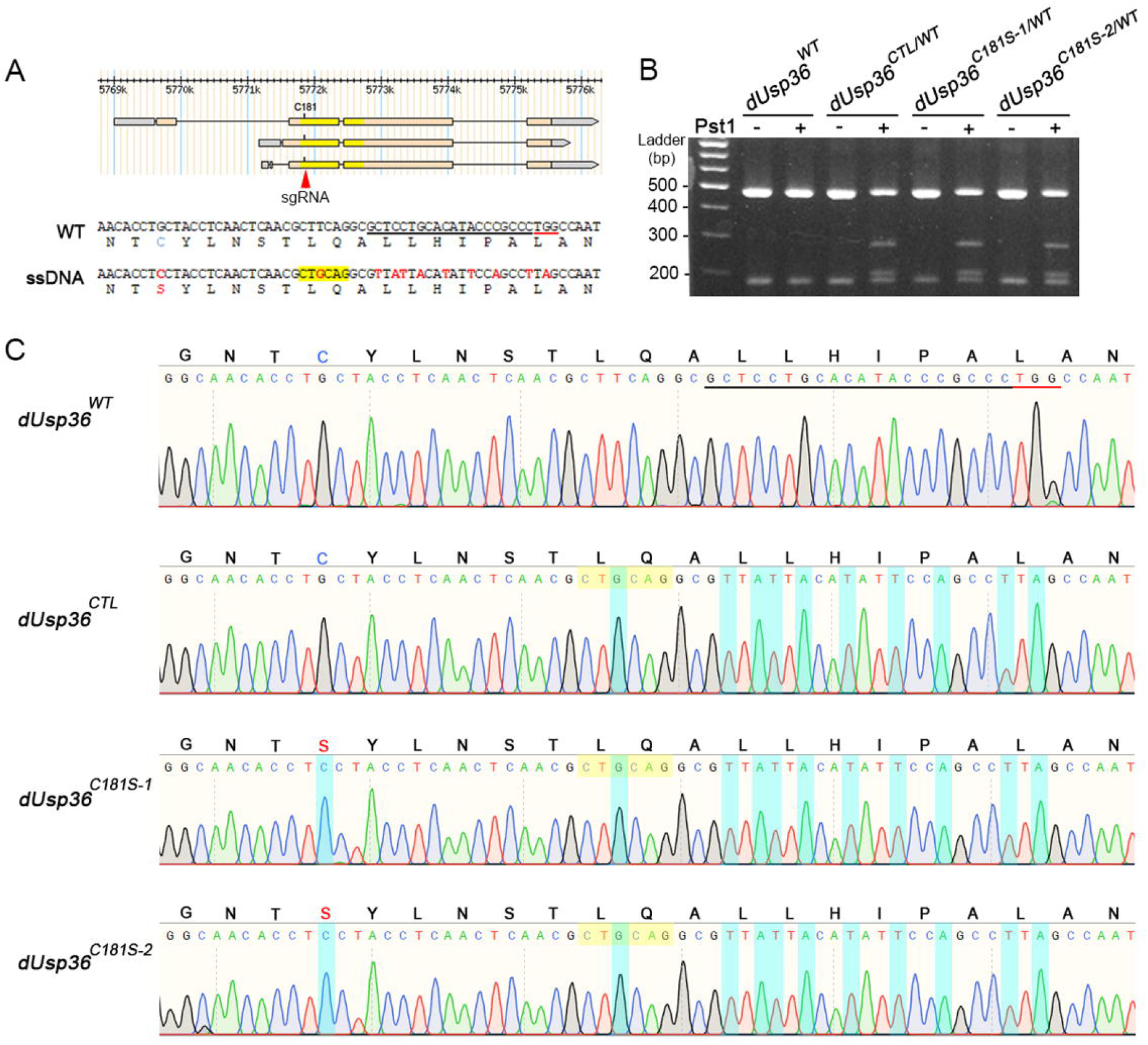
Generation of *dUsp36* catalytic mutants by CRISPR/Cas9 mutagenesis. (A) Schematic representation of the *dUsp36* locus with the three known isoforms. The position of the codon specifying the cysteine residue at position 181 is indicated as well as the gRNA (red triangle). WT: wild-type *dUsp36* sequence showing the targeted cysteine (in cyan), the sequence targeted by the gRNA (underlined in black) and the PAM (underlined in red). ssDNA: part of the ssODN sequence with the G-to-C transversion responsible for the C181S mutation and the silent mutations introduced for screening purposes (creation of a Pst1 restriction site and mutation of the PAM and of the gRNA to avoid re-cleaving of an edited chromosome). (B) Agarose gel showing that PCR products amplified from the *dUsp36*^*CTL*^, *dUsp36*^*C181S-1*^ and *dUsp36*^*C181S-2*^ alleles display a Pst1 restriction site which is absent of the wild-type allele. (D) Chromatograms showing the sequences obtained from the two independent *dUsp36* ^*C181S*^ alleles, as well as from the control allele, *dUsp36*^*CTL*^, which contains all the silent mutations introduced for screening purposes but not the G-to-C transversion responsible for the C181S mutation.

**Figure S2:**
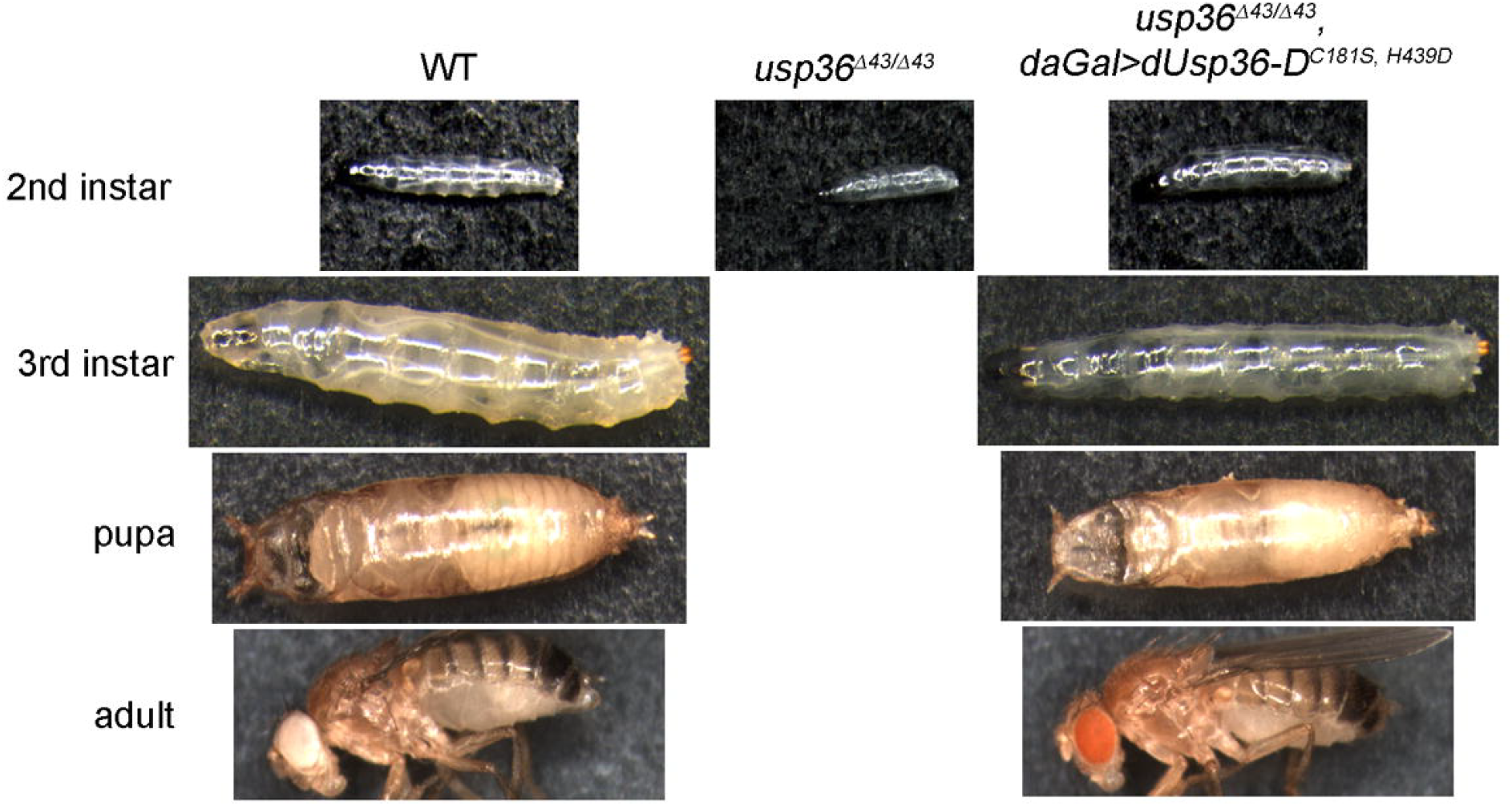
Rescue of the lethality of *dUsp36* null mutants by the ubiquitous expression of the double mutant dUSP36^C181S, H439D^ protein.

## Notes

### Competing Interest Statement

The authors have declared no competing interest.

### Summary of Updates

this version of the manuscript corresponds to the version submitted to Genetics

